# The Role of Conformational Dynamics in abacavir-Induced Hypersensitivity Syndrome

**DOI:** 10.1101/234260

**Authors:** James Fodor, Blake T. Riley, Itamar Kass, Ashley M. Buckle, Natalie A. Borg

## Abstract

Abacavir is an antiretroviral drug used to reduce human immunodeficiency virus (HIV) replication and decrease the risk of developing acquired immune deficiency syndrome (AIDS). However, its therapeutic value is diminished by the fact that it is associated with drug hypersensitivity reactions in up to 8% of treated patients. This hypersensitivity is strongly associated with patients carrying human leukocyte antigen (HLA)-B*57:01, but not patients carrying closely related alleles. Abacavir’s specificity to HLA-B*57:01 is attributed to its binding site within the peptide-binding cleft and subsequent influence of the repertoire of peptides that can bind HLA-B*57:01. To further our understanding of abacavir-induced hypersensitivity we used molecular dynamics (MD) to analyze the dynamics of three different peptides bound to HLA-B*57:01 in the presence and absence of abacavir or abacavir analogues. We found that abacavir and associated peptides bind to HLA-B*57:01 in a highly diverse range of conformations that are not apparent from static crystallographic snapshots. Further, the presence of abacavir has a direct impact on the dynamics and the conformational space available to peptides bound to HLA-B*57:01, likely influencing abacavir-induced immune self-reactivity. Our results support hypersensitivity models in which abacavir-binding alters the equilibrium proportions of neopeptide conformations in a manner favourable to TCR binding. Our findings highlight the need to also consider the role of dynamics in understanding drug-induced hypersensitivities at the molecular and mechanistic level. This additional insight can help inform the chemical modification of abacavir to prevent hypersensitivity reactions in HLA-B*57:01+ HIV patients whilst retaining potent antiretroviral activity.

## Introduction

Abacavir is an antiretroviral medication used for the treatment of human immunodeficiency virus (HIV) infection^1,2^. It is a prodrug that is converted by the liver^3^ to form the pharmacologically active compound carbovir 5′-triphosphate^4^, an analogue of guanosine that targets HIV reverse transcriptase. Abacavir has been found to elicit a drug hypersensitivity reaction in up to 8% of treated patients^5,6^, with hypersensitivity attributed to the prodrug itself^7,8^. Symptoms of abacavir hypersensitivity syndrome (AHS) include fever, malaise, nausea, diarrhoea and skin rash, and the condition can be fatal in severe cases^9^. AHS is strongly associated with patients carrying a human leukocyte antigen (HLA)-B*57:01 allele, and is mediated by the activation of HLA-B*57:01 restricted CD8+ T-cells^10-12^. In contrast, hypersensitivity is not observed in patients carrying the closely related alleles HLA-B*57:03 (Asp114Asn; Ser116Tyr), HLA-B*57:02 (Asp114Asn; Ser116Tyr; Leu156Arg), and HLA-B*58:01 (Met45Thr; Ala46Glu; Val97Arg; Val103Leu)^7,11^.

Mass spectrometry data characterizing the peptides eluted from HLA-B*57:01 cell lines treated with or without abacavir revealed abacavir alters the repertoire of peptides that are bound by HLA-B*57:01^7,8,13^. Crystallographic structures further revealed that abacavir binds non-covalently at the floor of the HLA-B*57:01 peptide-binding cleft, and makes contacts with F-pocket residues that differ in HLA-B*57:03, HLA-B*57:02 and HLA-B*58:01^7,8^. In doing so, abacavir changes the chemistry and shape of the peptide binding cleft, thereby altering the repertoire of peptides that can be presented by HLA-B*57:01.

To further our understanding of AHS and the interactions between HLA-B*57:01, bound peptide, and abacavir, we performed a series of MD simulations of three HLA-B*57:01-peptide complexes in the presence (PDB IDs 3UPR^8^, 3VRI and 3VRJ^7^) of abacavir, or with abacavir removed. We also performed MD simulations of three abacavir analogues (carbovir, didanosine, and guanosine) all bound to HLA-B*57:01, to serve as a comparison. These analogues are all chemically very similar to abacavir, but do not trigger a T-cell response^11^.

Our simulations show that abacavir can adopt a range of conformations in the HLA-B*57:01 antigen-binding cleft, as can the bound peptides. This suggests that conformational dynamics may play an underappreciated role in the recognition of HLA-B*57:01 by abacavir-specific T cell receptors (TCRs). We further show that the high levels of peptide dynamics, and partial dissociation of HLA-B*57:01-bound peptides may allow abacavir direct access to the antigen-binding cleft, and propose conformational dynamics to be a central tenet of all models of HLA-associated drug hypersensitivity. Overall our results suggest structures alone of the HLA-B*57:01-abacavir-peptide complexes are insufficient to account for the molecular basis of AHS.

## Results and Discussion

To compare the dynamics of HLA-B*57:01 +/-abacavir, or abacavir analogues, we utilized the three available abacavir-bound HLA-B*57:01 structures (PDB ID 3UPR, 3VRI and 3VRJ), and either removed abacavir or substituted an abacavir analogue into the location of abacavir, and subjected the structures to MD analysis. A summary of all the systems for which MD was performed is provided in Table 1 and the structural components are shown in Figure 1.

**Table 1.** Summary of MD simulations performed.

|  | Peptide |  |  |
| --- | --- | --- | --- |
|  | PDB ID: 3UPR | PDB ID: 3VRI | PDB ID: 3VRJ |
| **abacavir** | MHC + HSITYLLPV<br>+ abacavir | MHC + RVAQLEQVYI<br>+ abacavir | MHC + LTTKLTNTNI<br>+ abacavir |
| **carbovir** | MHC + HSITYLLPV<br>+ carbovir | MHC + RVAQLEQVYI<br>+ carbovir | MHC + LTTKLTNTNI<br>+ carbovir |
| **didanosine** | MHC + HSITYLLPV<br>+ didanosine | MHC + RVAQLEQVYI<br>+ didanosine | MHC + LTTKLTNTNI<br>+ didanosine |
| **guanosine** | MHC + HSITYLLPV<br>+ guanosine | MHC + RVAQLEQVYI<br>+ guanosine | MHC + LTTKLTNTNI<br>+ guanosine |
| **unbound** | MHC + HSITYLLPV | MHC + RVAQLEQVYI | MHC + LTTKLTNTNI |

**Figure 1:**
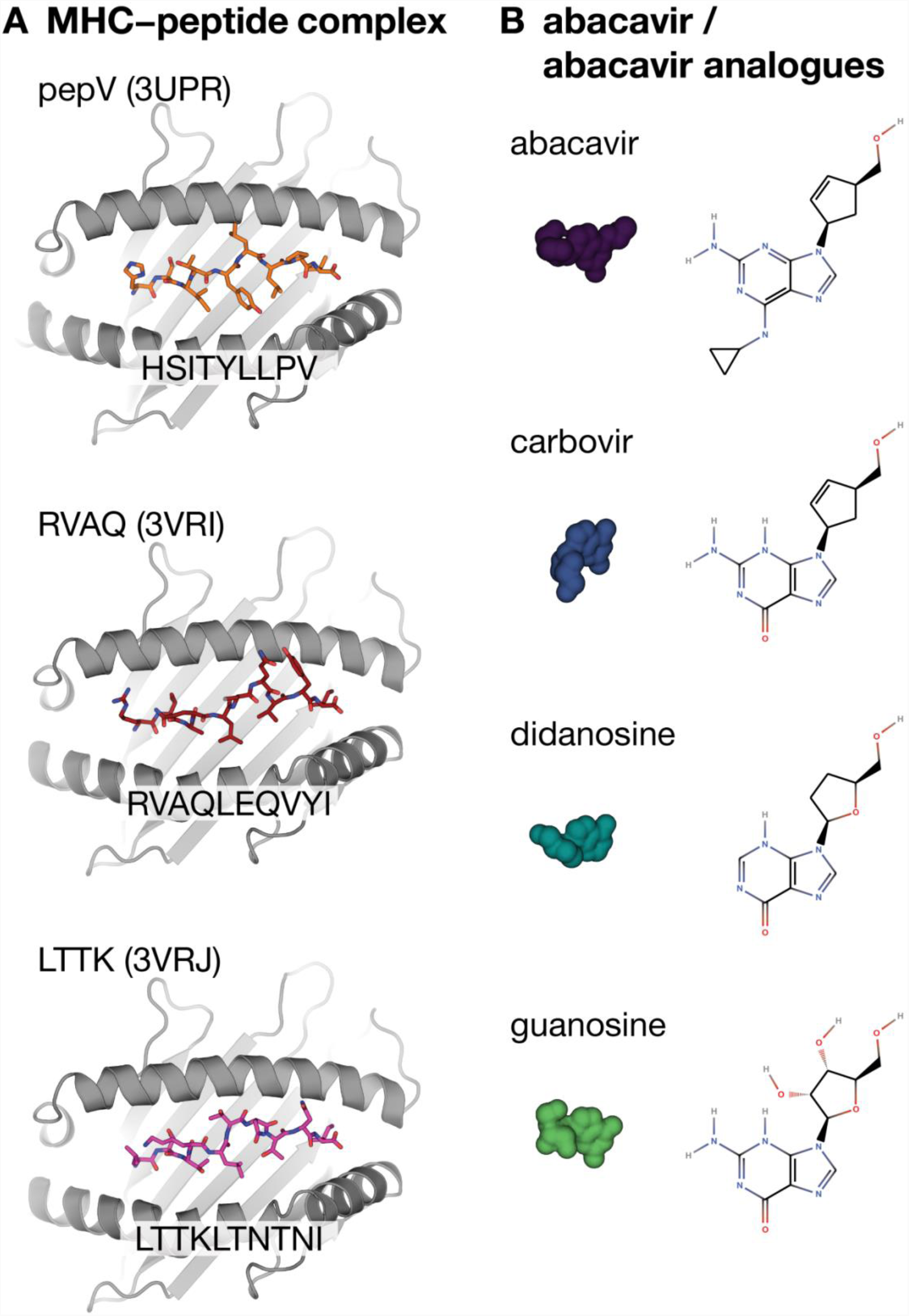
Overview of the structural components used to perform the MD simulations. (A) Structures of the HLA-B*57:01-peptide-abacavir complexes with abacavir removed. (B) Chemical structures and surface representations of abacavir and abacavir analogues.

### Peptides bound to HLA-B*57:01 in the presence of abacavir or abacavir analogues display conformational variation

To investigate the range of conformations adopted by the three HLA-B*57:01-bound peptides in the presence or absence of abacavir or abacavir analogues and over the course of the simulations, we defined three critical distances between residues in each peptide and residues in the underlying β-sheet (Figure 2A, C, E). The N- and C-terminal residues were selected to indicate the motion of each end of the peptide (denoted peptide edges), while the peptide residue showing the largest protuberance from the binding cleft (denoted peptide center) was also selected as a point of comparison. The distances between residues in each pair were then computed at each frame in the trajectory at 1 nanosecond intervals. The resulting distances are plotted in Figure 2.

**Figure 2:**
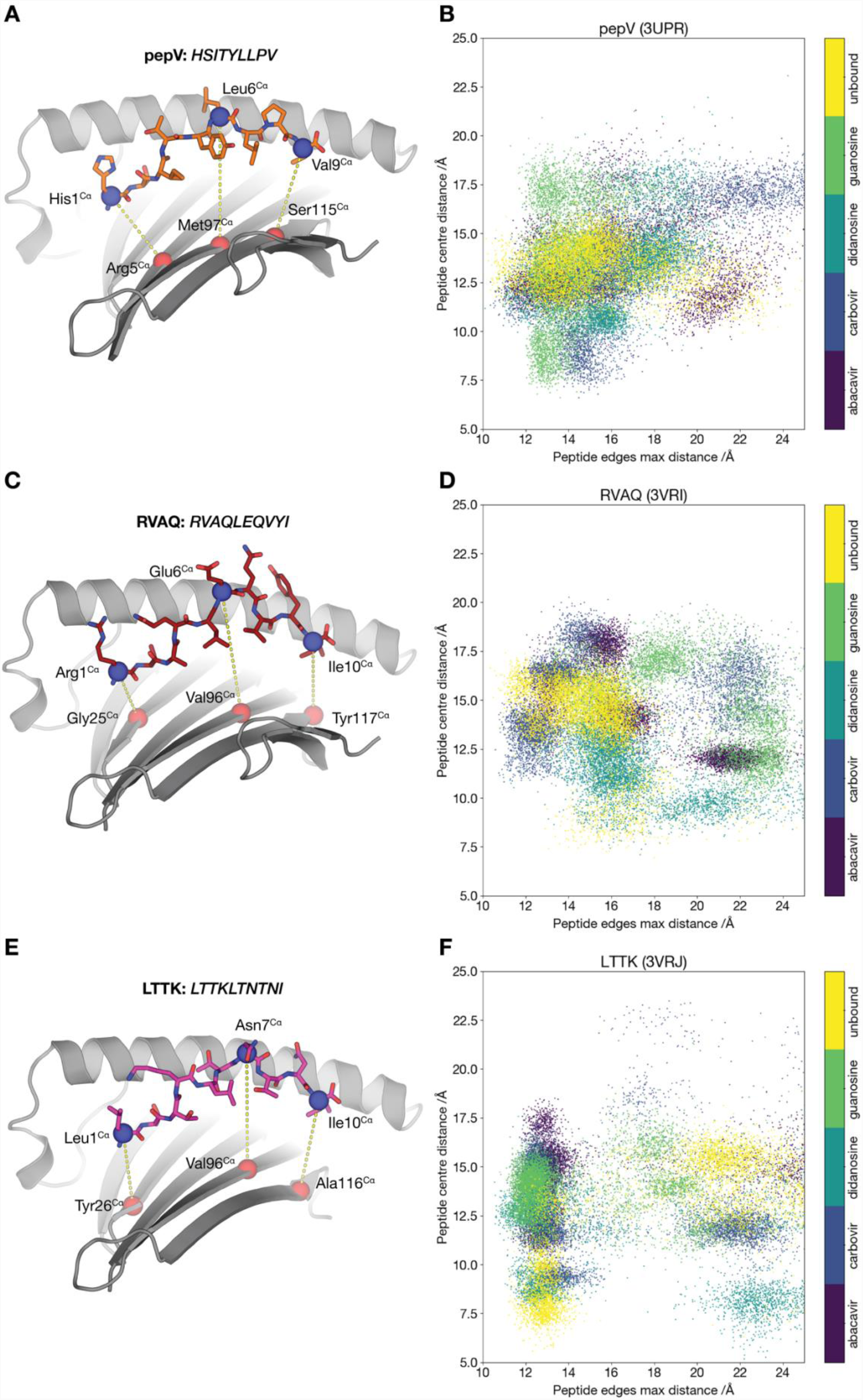
Cluster analysis of molecular simulation trajectories of all systems. (A, C, E) Residue pairs used for distance measurements. The MHC binding cleft is shown as a cartoon with the α2 helix omitted for clarity and the peptide shown in sticks. The Cα atoms of paired residues in the peptide and β-sheet are shown as spheres, coloured blue for peptide atoms and red for β-sheet atoms. The distance (in Å) is calculated between the atoms in each pair, shown as a yellow dotted line. (B, D, F) Plots of the distance between the Cα of the central peptide residue and the Cα of the underlying β-sheet residue (vertical axis), against the maximum of the distances between the Cα of the N- and C-terminal residues of the peptide and the Cα of the corresponding underlying β-sheet residue (horizontal axis). Each point on the graph corresponds to a single frame from each trajectory, with one frame shown for each nanosecond of simulation time. Each system is displayed in a different colour as per the key displayed with the figure.

Notwithstanding the static picture presented by the crystallographic structures of HLA-B*57:01-peptide-abacavir complexes, our simulations indicate that the complexes are in a state of constant conformational flux. The central residue of the peptide ranged in height above the underlying β-sheet from ∼ 7.5 to 20.5 Å in 3UPR, from 11 to 23 Å in 3VRI, and from 6 to 23 Å in 3VRJ (Figure 2). Likewise, the peptide edges ranged in height above the β-sheet from around 11 Å to up to 25 Å (Figure 2). Likewise, conformational flux was observed for the HLA-B*57:01-bound peptides in the absence of abacavir, and also when the abacavir analogues were present. Therefore, despite the similarity of the underlying systems, a wide range of conformations are adopted. Although most individual runs (three for each system) form a loose cluster on these graphs, there is no evidence of clear systematic differences between abacavir and any of its analogues, or between abacavir analogues and the unbound complex that differentiates the abacavir-containing systems.

This large conformational diversity can also be visualized directly by overlaying snapshots of the peptides from multiple frames in each simulation (Figure 3). High levels of conformational diversity can be observed, with substantial rearrangements of the peptide backbone within the antigen-binding cleft observed over the course of all the simulations. As before, there are no features distinguishing abacavir-bound simulations from the other systems.

**Figure 3:**
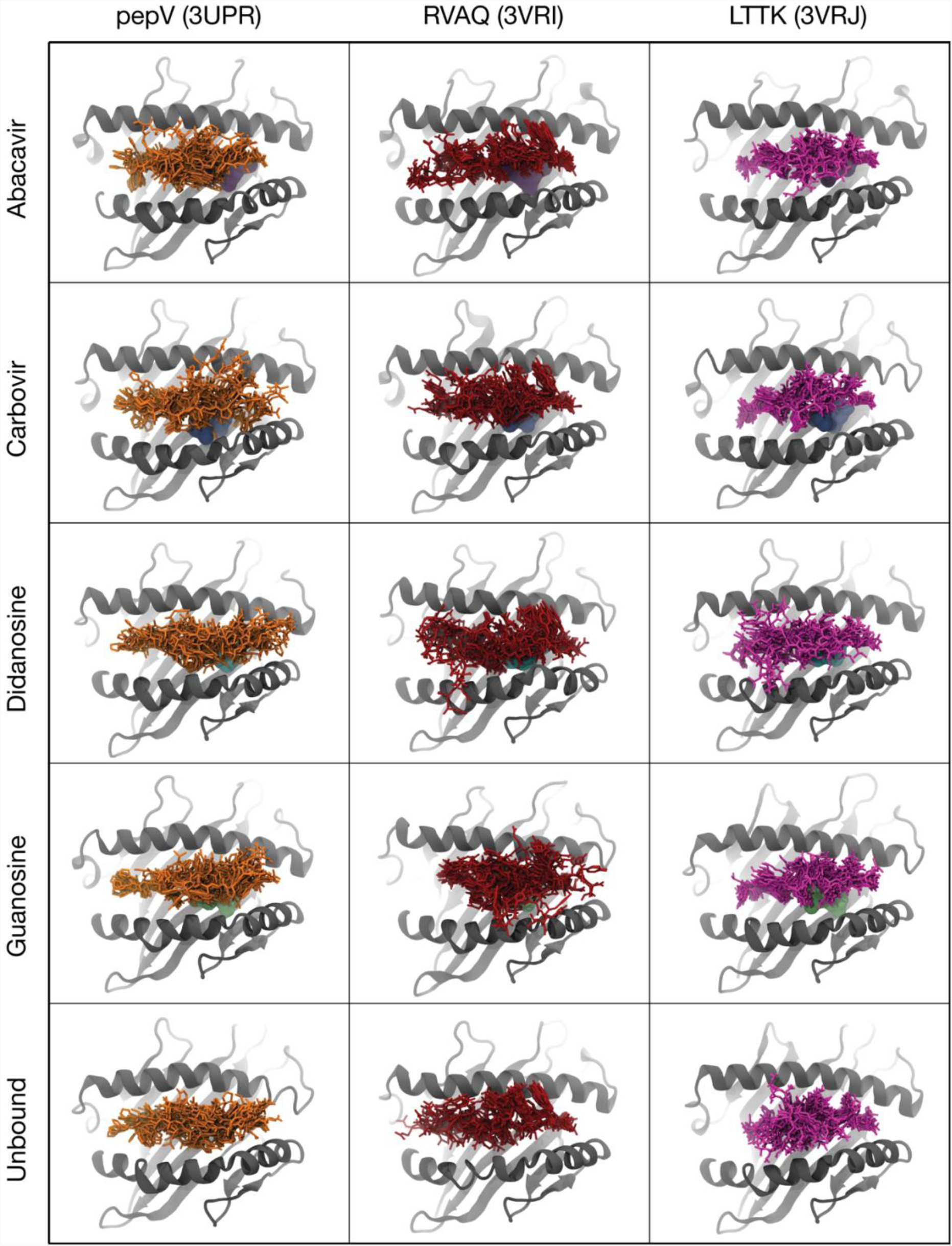
Conformational heterogeneity of HLA-B*5701-bound peptides. Overlay of peptide conformations from 10 frames of each simulation of structures of unbound HLA-B*57:01 and HLA-B*57:01 bound to abacavir or abacavir analogues. The peptide is shown in sticks (pepV in orange, RVAQ in brick red, LTTK in magenta), HLA-B*57:01 in gray cartoon, and the compound (if present) shown as a surface representation.

### HLA-B*57:01-bound peptides partially detach from the peptide binding groove

As a further measure of structural variability, root mean squared deviation (RMSF) relative to starting positions was calculated for all residues in each system over the course of the trajectory, and the results for the peptide residues plotted in Figure 4. The majority of residues for most systems show relatively high RMSFs of 3 Å or more, with some of the terminal regions having very high RMSFs of up to 8 Å. This is consistent with the observation that both the N- and C-termini of the peptide escape from the MHC antigen-binding cleft in several of the simulations (Table 2). Similar results were obtained in previous, shorter MD simulations, in which the C-terminus of a range of self-peptides detached from the peptide binding groove of HLA-B*57:01 in the absence of abacavir^14^. However, abacavir does not show any systematic tendency to either increase or decrease peptide RMSF relative to either the other analogues or to the unbound system. Overall these results clearly indicate high levels of peptide dynamics over the course of the simulation, but no distinctive effects of abacavir which differentiates it from the other systems are evident.

**Table 2:**
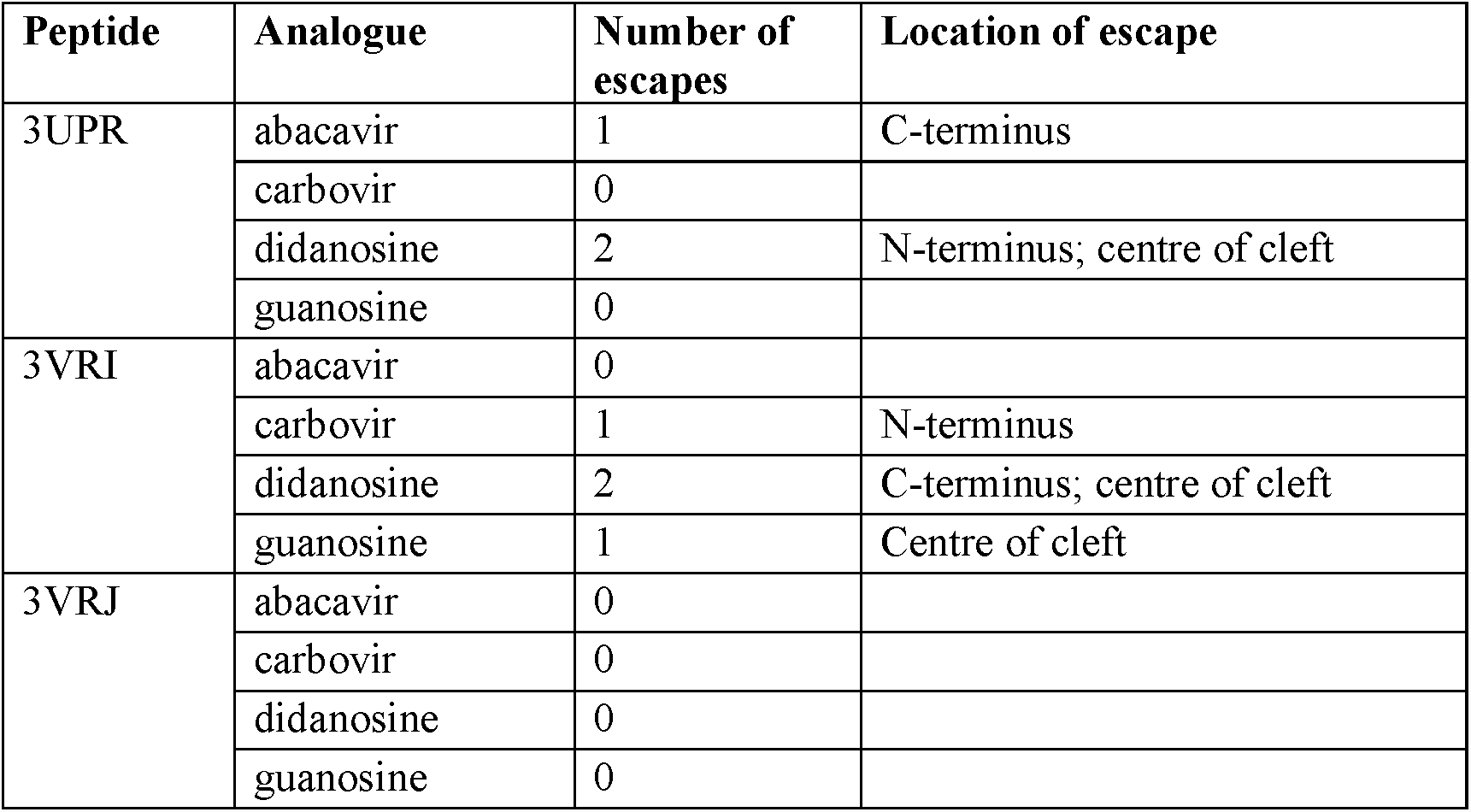
Number of abacavir or abacavir-analogue escapes from the antigen-binding cleft.

**Figure 4:**
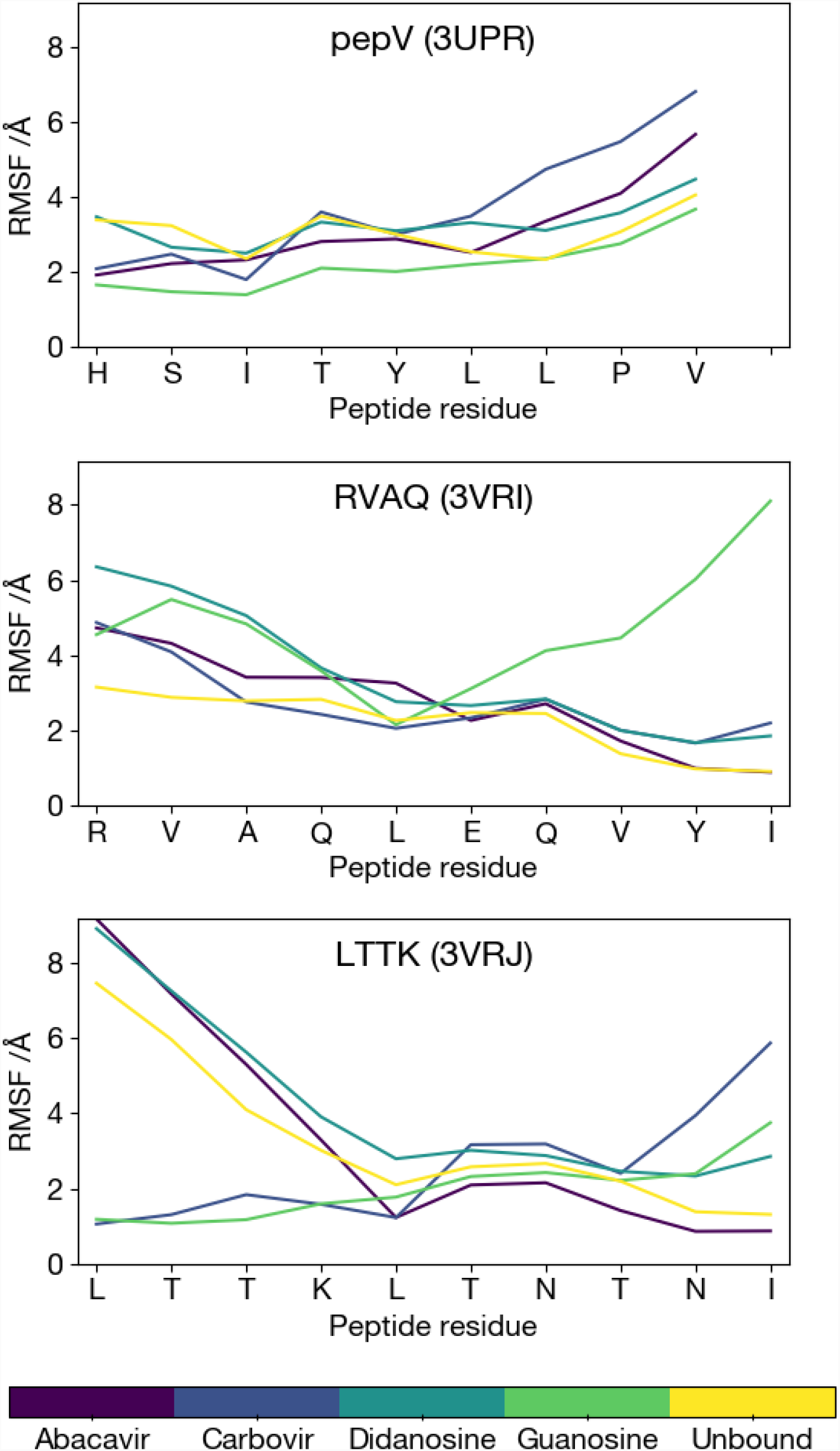
Peptide dynamics within HLA-B*57:01 during the simulations. Root mean squared fluctuation (RMSF) for each residue of the peptide for 3UPR (top), 3VRI (middle), and 3VRJ (bottom) with or without abacavir or abacavir analogues.

### Abacavir and abacavir analogues can escape the HLA-B*57:01 antigen-binding cleft

Examination of snapshots taken throughout the simulations of abacavir in the HLA-B*57:01 antigen-binding groove show that abacavir can bind to HLA-B*57:01 in many similar but distinct conformations. In some cases abacavir and its analogues escape from the binding cleft (Table 2), with escapes occurring as a result of partial detachment of the peptide at the N- and C-termini, and also from the centre of the binding cleft without any partial detachment of the peptide. One possible explanation for these results is that abacavir and the abacavir analogues do not bind very stably to the MHC-peptide complex. For the abacavir analogues this is plausible given they were modeled into the HLA-B*57:01 structures and, unlike abacavir, have not been proven to bind HLA-B*57:01. This explanation, however, only accounts for the pattern observed for the 3VRI systems, and not for the absence of any escapes in the 3VRJ system, or the fact that abacavir escapes in the 3UPR system while carbovir and guanosine do not. Overall it is difficult to provide a clear rationale for these results in terms of single structural differences. Indeed, it seems likely that all of the abacavir analogues would escape from the binding clefts if the simulations could be run for long enough timescales, and that the observed differences may be due largely to the inherently stochastic nature of MD.

### Abacavir and abacavir analogues can bind to HLA-B*57:01 in a range of conformations

Examination of overlays of snapshots from the MD trajectories shows that abacavir exhibits a considerable range of conformations within the HLA-B*57:01 antigen-binding cleft (Figure 5). Particularly common are rotations of the cyclopentyl and cyclopropyl moieties, as well as translational motion of the entire compound within the binding cleft. In all cases it is evident that the conformation observed in the original crystallographic structure is but one among a large conformational ensemble.

**Figure 5:**
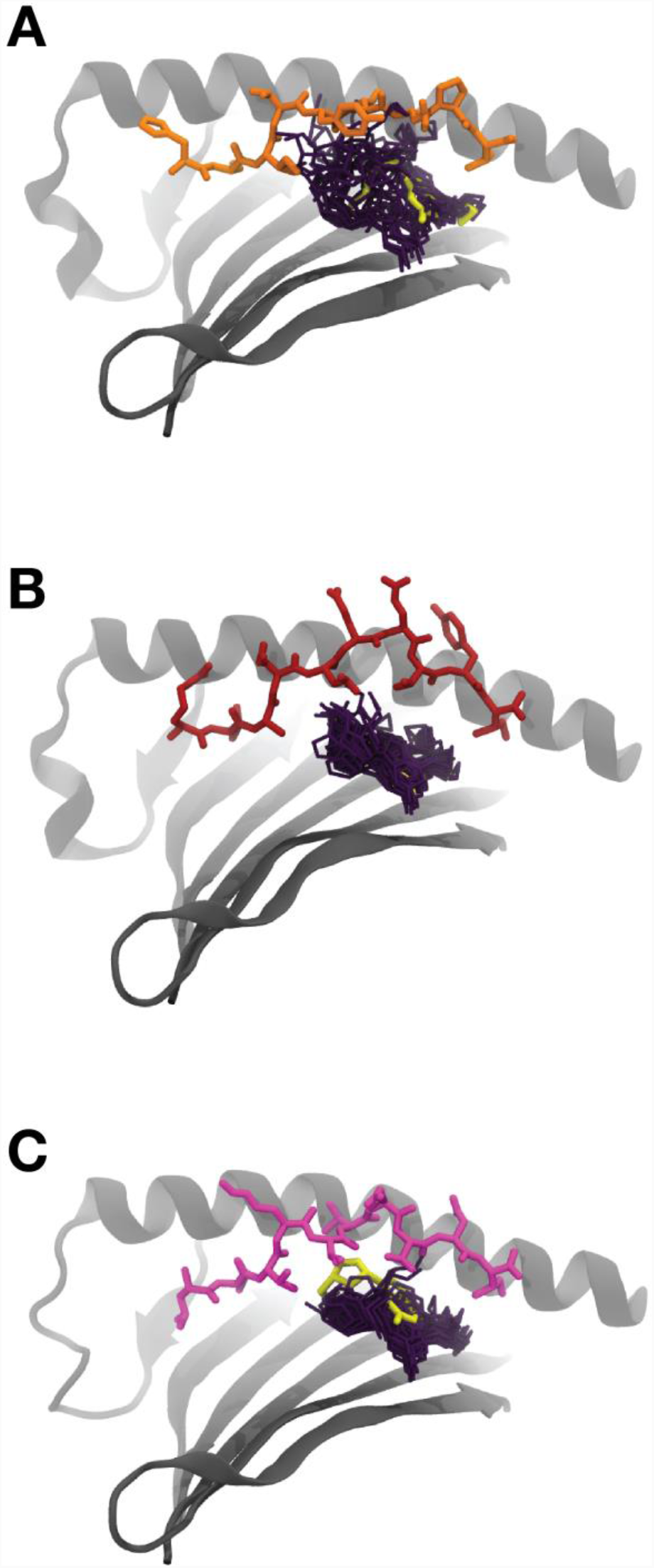
Conformational heterogeneity of abacavir. Overlay of abacavir conformations from 10 frames of each simulation on a static HLA-B*57:01 and peptide for A) 3UPR, B) 3VRI, and C) 3VRJ. The peptide is shown in sticks (pepV in orange, RVAQ in brick red, LTTK in magenta), HLA-B*57:01 in grey cartoon with the α2 helix omitted for clarity, and abacavir in deep purple sticks. The conformation of abacavir as observed in the corresponding crystallographic structure is also shown in yellow sticks.

Our observation that abacavir retains considerable flexibility/mobility while lodged between HLA-B*57:01 and the bound peptide implies that the loss of entropy upon HLA-B*57:01-peptide-abacavir complex formation is smaller than would be the case were abacavir to remain totally rigid, thus favouring complexation. The flexibility of abacavir may explain its specificity toward HLA-B*57:01. As previously noted, closely related alleles such as HLA-B*57:03, HLA-B*57:02, and HLA-B*58:01 do not bind abacavir and thus are not influenced by it^7,11^. These alleles differ from HLA-B*57:01 at a small number of sites with bulky amino acids, mainly at positions Ala46, Val97, Ser116 and Leu156. During simulations, abacavir is observed to interact with these residues. As a consequence, mutations in those residues that introduce a bulkier amino acid, for example Ser116Tyr or Val97Arg, likely create steric clashes which reduce the conformational space available to abacavir when bound to these alleles. As we have shown that abacavir influences complex stability via its effect on the shape of the binding groove, rather than through direct interactions with binding pockets, bulkier residues may destabilize the pMHC-abacavir complex by limiting these effects.

### Abacavir is highly dynamic when lodged between HLA-B*57:01 and the bound peptide

Our results strongly indicate that the nature and causes of abacavir’s hypersensitivity reaction cannot be explained solely through analysis of static structures. Abacavir and the peptides with which it binds vary in conformation to such an extent that any analysis based on the presence or absence of particular steric interactions or hydrogen bonds will almost inevitably fail to identify features which are consistently present in abacavir and only abacavir. Further, despite performing a combined total of 90 microseconds of MD, no clear structural features or interactions were found to be present in abacavir systems but not any of its analogues, nor were there any such systematic differences between the abacavir analogues as a group and the system with only the peptide.

One outstanding question is whether the three peptides considered in the crystallographic studies bind to HLA-B*57:01 in the absence of abacavir, and also whether they bind in the presence of any of the abacavir analogues. While previous work has indicated that such binding has not been observed^7,8^, our simulation results fail to indicate that the MHC-peptide-abacavir complex is any more stable than the MHC-peptide or MHC-peptide-analogue complexes. Indeed, our results show that all of these systems are highly dynamic, with one or other peptide termini commonly becoming dislodged from the binding cleft, and abacavir or one of its analogues escaping the binding cleft entirely in many of the simulations. There are also other reports of peptide escape allowing access to drugs, for example in allopurinol hypersensitivity^15^, and also other cases of partial peptide detachment reported for various peptides bound to HLA-B*57:01^14^ and HLA-B*44:02^16^. These findings indicate that in all likelihood none of the MHC-peptide-abacavir or MHC-peptide-analogue complexes are especially stable, and differ largely in terms of their relative association and dissociation constants.

Previous studies have focused on the interaction of the abacavir cyclopropyl group with the Val97 and Ser116 residues unique to HLA-B*57:01 as explaining why only abacavir (which alone of its analogues possesses this cyclopropyl group) and only HLA-B*57:01 (which alone of its similar alleles possesses this residue pair) elicit recognition by T cells^7,8^. However, our simulations indicate that abacavir does not adopt a consistent conformation with respect to these residues (Figure 6). Conversely, similar forms of interaction are also observed to occur with the abacavir analogues. As such, even if some of these interactions are important for eliciting T cell recognition by abacavir, this can only be a matter of degree, and not due to the binary existence or non-existence of certain interactions.

**Figure 6:**
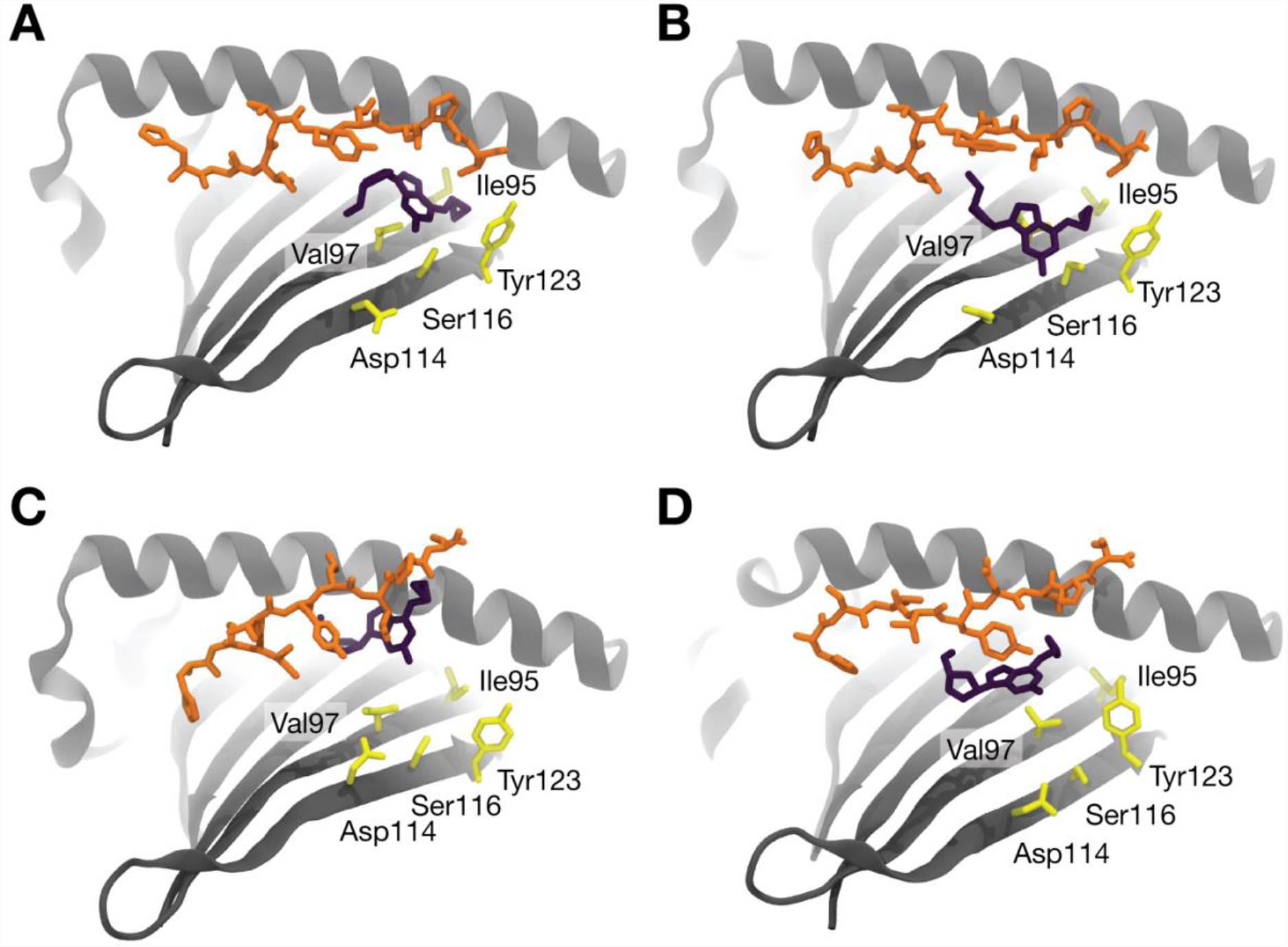
Key HLA-B*57:01 residues and their conformation relative to that of abacavir. HLA-B*57:01 is shown in gray cartoon with residues Ile 95, Val97, Asp114, Ser116, and Tyr123 shown in yellow sticks. Abacavir is shown in deep purple sticks and the pepV peptide pepV in orange sticks. The MHC α2 helix has been omitted for clarity. (A) The starting frame from run 1, and a conformation from each of the independent replicates (B) run1, (C) run2, and (D) run3 are shown, illustrating how abacavir does not maintain a consistent position or orientation with respect to these residues.

AHS is attributed to the altered repertoire model in which abacavir non-covalently interacts with the binding-cleft of HLA-B*57:01, altering the repertoire of peptides that can bind and allowing the presentation of neoantigens, preferring Ile, Leu, Val or Ala at PΩ, that are considered foreign to circulating CD8+ T cells^7,8,13^. Our MD data suggests HLA-B*57:01-bound peptides bearing abacavir-compatible PΩ residues adopt a highly diverse range of conformations when abacavir is bound, and that these altered conformations also play a role in abacavir-induced T cell activation. The intricacy of the short- and long-lived interactions between the TCR and HLA-B*57:01-peptide is consistent with the role of conformational dynamics in the activation of the immune system^17-23^. The TCR scans a wide range of conformations in each system existing in a conformational ensemble. Glimpses of such scanning have been reported, for example flexibility within crystal structures of B*3508-LPEP in complex with SB27 TCR, and subsequent MD studies suggested a scanning motion of the TCR on top of the pMHC^21,24^. Some particular conformations will facilitate binding to a higher degree than others, and if these conformations are more common in the abacavir complex than in the other putative complexes, then in equilibrium the TCR will be bound to the abacavir complexes more frequently, thereby eliciting an immune response. Much longer MD simulations, and possibly more advanced sampling approaches, would be necessary to fully characterize the relative populations of different conformations in each ensemble, and thereby determine precisely which conformations are responsible for any preferential TCR response to abacavir.

Our MD simulations also indicate HLA-B*57:01-bound peptides can partially dissociate out of the peptide-binding cleft, allowing abacavir direct access. Partial peptide detachment has similarly been attributed to the exposure of the allopurinol/oxypurinol binding site within the cleft of HLA-B*58:01, which is associated with HLA-associated drug hypersensitivity^15^. This hypersensitivity, described by the p-i (pharmacological interaction with immune receptor) model, is due to the direct, reversible binding of the drug to the HLA-B*58:01 antigen-binding cleft which stabilizes novel peptide conformations that immediately stimulate a T cell response. Given the partial peptide detachment observed in our simulations it is possible that, in addition to the altered repertoire model, that the p-i model may also be a driver for AHS. This ‘dual’ model would account for the observed immediate activation of abacavir-reactive T cells^25^ in addition to the delayed abacavir-induced T cell response characteristic of a dependency on conventional antigen-processing pathways^7,11^. Notably, drug-induced novel peptide conformations are central to both models.

Finally, our results constitute strong evidence against static structure explanations of abacavir-induced hypersensitivity which focus on binary interactions between MHC-peptide-abacavir and a T cell response stimulated by a single-conformation peptide, but rather support a mechanism involving selection from amongst dynamical pMHC conformational ensembles.

## Concluding remarks

Our MD simulations highlight that HLA-B*57:01-peptide-abacavir crystal structures, which represent static molecular snapshots, do not fully provide an explanation for the strong association of abacavir hypersensitivity with HLA-B*57:01. We show that whether or not abacavir or any of its analogue compounds are present in the HLA-B*57:01 antigen-binding cleft, the peptides exhibit high levels of conformational flexibility. Our MD data does not support the notion that any particular differences in specific conformations or degree of flexibility can account for abacavir’s unique immunogenicity. Instead, we propose that hypersensitivity is the result of abacavir altering the equilibrium proportions of different peptide conformations in a manner favourable to TCR binding. Further, we propose the high levels of peptide dynamics, and partial dissociation of HLA-B*57:01-bound peptides, may also allow abacavir direct access to the antigen-binding cleft. Based on our observations it is possible that AHS is driven by both the p-i and altered repertoire hypersensitivity model. We propose that conformational dynamics is a central tenet of all models of HLA-associated drug hypersensitivity, and should be characterized so as to complement crystallographic studies and *in silico* docking approaches^26^. The integration of such techniques, alongside the cellular studies and genetic associations reported to drugs, provides for a more comprehensive molecular and mechanistic insights into HLA-associated drug hypersensitivities. Given HLA-associated adverse drug reactions have been reported for over 10 drugs, ranging from anticonvulsants to antibiotics (see ^27^ for a review), this united approach may accelerate the development of safer and more effective drugs of clinical use for a range of human health conditions.

## Materials and Methods

### Computational resources

Parameterization of abacavir analogues was performed on Multi-modal Australian ScienceS Imaging and Visualisation Environment (MASSIVE), while MD simulations were performed on in-house hardware (NVIDIA TITAN X Pascal GPU).

### Systems preparation

The coordinates for starting structures of peptide-MHC complexes with abacavir were taken from PDB IDs 3UPR^8^, 3VRI and 3VRJ^7^, which contain bound-peptides HSITYLLPV (pep-V), RVAQLEQVYI (RVAQ), and LTTKLTNTNI (LTTK), respectively. For each unique pMHC complex, systems with and without abacavir were simulated. Initial abacavir geometry was extracted from a crystal structure (PDB ID 3UPR^8^), while structures for abacavir analogues carbovir, guanosine, and didanosine were obtained from PubChem^28^, and positioned in the binding cleft by aligning them with the carbon and nitrogen atoms of the central purine moiety of abacavir using PyMOL. Each protein, with protonation states appropriate for pH 7.0 as determined by PROPKA^29,30^, was placed in a rectangular box with a border of at least 10 Å, explicitly solvated with TIP3P water^31^, sodium counter-ions added, and parameterized using the AMBER ff99SB all-atom force field^32-34^. After an energy minimization stage consisting of at least 10,000 steps, an equilibration protocol was followed in which harmonic positional restraints of 10 kcal Å^2^ mol^-1^ were applied to the protein backbone atoms. The temperature was incrementally increased while keeping volume constant from 0 K to 300 K over the course of 0.5 ns, with Langevin temperature coupling relaxation times of 0.5 ps. After the target temperature was reached, pressure was equilibrated to 1 atm over a further 0.5 ps using the Berendsen algorithm^35^.

### Abacavir analogues parameterisation

For each of the abacavir analogues, Gaussian 09^36^ was used to optimize its conformation and to calculate its molecular electrostatic potential (ESP). This was done at the HF/6-31++G* level of theory, to allow for polarization effects. The ANTECHAMBER module of AmberTools 12^37^ was then used to generate the force field libraries by assigning RESP-fitted charges, as well as Generalized Amber Force Field (GAFF)-derived parameters and atom types^38^ for use in Molecular Dynamics (MD) simulations.

### Molecular Dynamics (MD)

Following equilibration and parameterization, production MD simulation runs were performed in the NPT ensemble using periodic boundary conditions and a time step of 2 fs. Temperature was maintained at 300 K using the Langevin thermostat with a collision frequency of 2 ps, and electrostatic interactions computed using an 8 Å cutoff radius and the Particle Mesh Ewald method^39^. Pressure was maintained at a constant 1 atm using Berendsen pressure bath coupling with a time coefficient of 0.1 ps^35^. Each simulated system was repeated three times from the same starting structure but with different starting velocities, with each run extending for 2 microseconds.

### MD analyses

After the simulations were completed, the three trajectories of each system were concatenated for analysis. Root Mean Square Fluctuation (RMSF) analyses and atomic distance computations were performed using VMD version 1.9.4^40^. Images were rendered using PyMOL^41^. RMSD and RMSF of backbone heavy atoms with respect to their initial structure were calculated every 1 ns, after performing a least-squares fit^42^ to the initial structure. RMSF results were reported as the average RMSF per residue backbone throughout simulations. Clustering analysis was performed using python.

## Acknowledgments

NAB is funded by an Australian Research Council (ARC) Future Fellowship (110100223). This work was supported by the Victorian Life Science Computational Initiative. The authors wish to thank Dr. Grischa Meyer for providing helpful scripts, as well as the Monash eResearch Centre.

## Author Contributions

AMB and NAB designed the study. JF and IK performed molecular dynamics simulations. JF performed analysis of the MD trajectories. BTR contributed to analysis and prepared the figures. IK, JF, BTR, AMB and NAB wrote the manuscript.

